# RNase R, a new virulence determinant of *Streptococcus pneumoniae*

**DOI:** 10.1101/2021.02.24.432507

**Authors:** Cátia Bárria, Dalila Mil-Homens, Arsénio M. Fialho, Cecília Maria Arraiano, Susana Domingues

## Abstract

Pneumococcal infections have increasingly high mortality rates despite the availability of vaccines and antibiotics. The increase of bacterial resistance to antibiotics urges the discovery of new alternative therapeutics. Therefore, the identification of new virulence determinants, and the understanding of the molecular mechanisms behind pathogenesis and pneumococcal-host interactions has become of paramount importance in the search of new targets for drug development. The exoribonuclease RNase R has been involved in virulence in a growing number of pathogens. In this work, we have used *Galleria mellonella* as an infection model to demonstrate that the presence of ribonuclease R increases the pneumococcus virulence. Although the absence of RNase R does not affect exponential growth, the ability of the RNase R deleted strain to replicate in the hemolymph is compromised. Larvae infected with the RNase R mutant strain show an increased expression level of antimicrobial peptides, and have a lower bacterial load in the haemolymph in the later stages of infection, leading to a higher survival rate. Interestingly RNase R carrying pneumococci suffer a sudden drop in bacterial numbers immediately after infection, resembling the eclipse phase observed after intravenous inoculation in mice. Together our results suggest that RNase R might be involved in the ability of pneumococci to evade the host immune response, probably by interfering with internalisation and/or replication inside the larval hemocytes.

## INTRODUCTION

*Streptococcus pneumoniae*, is an opportunistic pathogen that can be usually found as a harmless commensal of the human upper respiratory tract. Although this bacterium can colonize the nasopharynx of healthy individuals in an asymptomatic manner, propagation of pneumococcal cells beyond its niche along the nasal epithelium can lead to invasive diseases (reviewed in [1]). Dissemination either by aspiration, bacteraemia or local spread, can result in infections, such as pneumonia, meningitis, otitis media and septicaemia. In fact, in 2017 *S. pneumoniae* was included by the World Health Organization (WHO) as one of the 12 priority pathogens. The switch that makes this bacterium change from colonizer to pathogen is triggered by the opportunity to invade the bloodstream, tissues or organs of the host. Translocation from the nasopharynx to deeper tissues exposes *S. pneumoniae* to different environmental niches, which require a rearrangement of the bacterial cell. We have recently demonstrated that ribonuclease R (RNase R) affects translation and overall decreases protein synthesis in *S. pneumoniae* [2]. Appropriate translation is essential for the modulation of gene expression needed to cope with environmental changes, including favourable conditions to cause infection, which triggers the expression of virulence genes. A comprehensive understanding of the virulence mechanisms is of extreme importance for the development of new strategies to combat this bacterium. RNase R is important for virulence in several pathogens being upregulated in stress conditions [3] and we have shown that it is upregulated under cold shock in *S. pneumoniae* [4]. RNase R belongs to the RNase II/RNB family of enzymes, which is present in all domains of life [5]. This enzyme degrades RNA in the 3’-5’ direction in a processive and sequence-independent manner, being the only 3’-5’ exoribonuclease able to degrade highly structured RNAs [5]. RNase R is the only member of the RNB family present in *S. pneumoniae*, suggesting a major role in this microorganism. Therefore, we wanted to test if the absence of RNase R could compromise *S. pneumoniae* ability to cause infection.

Different *in vivo* model systems exist to study *S. pneumoniae* interaction with the host [6–8] but they all suffer from the issues of practicality, ethics and cost associated with mammalian hosts. To overcome these limitations, alternative infection model organisms have been investigated, such as the larvae of the wax moth *Galleria mellonella*. *G. mellonella* larvae have been proven to be an effective model host for infection studies in several microorganisms ([9-12] reviewed by [13]) including *S. pneumoniae* [14]. Additionally, larvae can be reared at temperatures ranging from 20°C to 30°C and the infection studies can be conducted between 15°C to above 37°C, which enables experiments that attempt to mimic a human environment [15, 16]. The innate immune system of *G. mellonella* involves cellular and humoral immune responses. The cellular response is mediated by phagocytic cells, termed hemocytes, that are involved in phagocytosis, encapsulation and clotting. Hemocytes are found within the hemolymph and function analogously to mammalian neutrophils in terms of their ability to phagocytose and kill pathogens, through the production of superoxides [17–20]. The humoral response is composed by soluble effector molecules which immobilize and/or kill the pathogen when released into the hemolymph. This response include the induced expression of antimicrobial peptides, as well as several plasma proteins that serve as opsonins, complement-like proteins, and release of the phenol oxidase system that culminates in the synthesis of the dark pigment melanin [21–23] reviewed by [13]).

In this work we have used *G. mellonella* to study the impact of the exoribonuclease RNase R in the pneumococcus virulence. We show that RNase R has an important role in the pathogenicity of *S. pneumonia* and present evidences suggesting that this enzyme might have a role in host/microbe interaction. Our results suggest that RNase R might promote the pneumococcus ability to evade the immune response of the host, probably by interfering with the internalization in the hemocytes.

## EXPERIMENTAL PROCEDURES

### Bacterial Strains, Plasmids, Insects and Growth Conditions

All bacterial strains and plasmids used in this study are listed in Table SI (Supplementary Material). *S. pneumoniae* strains are isogenic derivatives of the JNR7/87 capsulated strain – TIGR4. *S. pneumoniae* strains were grown in Todd Hewitt medium supplemented with 0.5% yeast extract (THY), at 37°C without aeration or in THY agar medium supplemented with 5% sheep blood (Thermo Scientific) at 37°C in a 5% CO_2_ atmosphere. When required growth medium was supplemented with 3 μg/ml chloramphenicol (Cm) or 1 μg/ml erythromycin (Ery).

*G. mellonella* larvae were reared on their natural food, beeswax, and pollen grains at 25°C in darkness prior to use. Larvae weighing 250 ± 25 mg were used.

### Flow Cytometry Analysis

Cellular growth was evaluated using S3e Cell Sorting equipment (Biorad). Overnight cultures of *S. pneumoniae* TIGR4 wild type and *rnr* mutant strain were diluted in pre-warmed THY to a final OD_600_ of 0.05 and incubated at 37°C. At OD_600_ ≈ 0.3 a volume of 15 mL of culture was centrifugated at 3800 rpm for 10 min. The cellular pellet was resuspended in sterile PBS with propidium iodide (PI) at a final concentration of 10 μg/mL and incubated 10 min in dark. Cells were centrifugated at 823 × *g* for 10 min and subsequently resuspended in 1 mL sterile PBS. 20 μL of the cell resuspension were transferred into a flow cytometry tube containing 1 mL of sterile PBS and analysed in the flow cytometry equipment. Fluorescence was detected using FL1 and FL2 filters. Live cells and death cells from old cultures were used as controls to set up gating.

### *Galleria mellonella* Killing Assay

For *G. mellonella* infection, overnight cultures of *S. pneumoniae* TIGR4 wild type and derivatives were diluted in pre-warmed THY to a final OD_600_ of 0.05 and incubated at 37°C. At OD_600_ ≈ 0.3 an appropriate volume was collected to assure that the same number of cells was used for all the strains. Cells were then harvested by centrifugation and resuspended in sterile PBS in a series of 10-fold serial dilutions corresponding to the number of 5×10^6^ bacterial cells per volume of injection. A micrometer was adapted to control the volume of a microsyringe and 5 μL aliquots of each dilution were injected into *G. mellonella*, via the hindmost left proleg, which had been previously surface sterilized with 70% ethanol. Control larvae were injected with the same volume of sterile PBS. Following injection, larvae were placed in glass petri dishes and incubated in the dark at 37°C for 3 days. For each condition, the survival and appearance of 10 larvae were followed at 24-h intervals. Larvae were considered dead when they displayed no movement in response to touch.

### *G. mellonella* RNA extraction

Sets of 20 larvae were infected with *S. pneumoniae* strains at a concentration of 5×10^6^ bacterial cells per larvae, as previously described for the killing assays. At 1, 6, 12, and 18 h after injection, three living larvae per set were cryopreserved, sliced, and homogenized in 1 ml of TRIzol reagent (Sigma-Aldrich). Whole-animal RNA was extracted according to the manufacturer’s protocol. After extraction, RNA was treated with an RNase-free DNase set (Qiagen). The purified RNA was quantified spectrophotometrically (NanoDrop ND-1000).

### Quantitative real-time PCR

The transcriptional levels of genes encoding the *G. mellonella* antimicrobial peptides gallerimycin, galliomycin, inducible metalloproteinase inhibitor (IMPI), and lysozyme were determined with a Real Time Thermal cycler qTower (Analytical Jena) system using a SensiFast SYBR kit (Bioline) according to the supplier’s instructions. cDNA was synthesized from 1 μg of purified RNA with a SensiFast cDNA synthesis kit (Bioline). The primers used are listed in Table S2 (Supplementary Material). All samples were analyzed at least in triplicate, from three independent biological samples. The amount of mRNA detected was normalized using the mRNA value of the housekeeping gene actin. Relative quantification of gene expression was calculated by using the ΔΔCT (CT is threshold cycle) method [24].

### Extraction of larval hemolymph

To determine the number of viable bacteria in the hemocoel during the course of infection at 1, 6 and 18 h after injection, the hemolymph of infected larvae was collected as previously described [12]. Briefly, three living larvae were anesthetized on ice and surface sterilized with ethanol. The larvae were bled, and the outflowing hemolymph was immediately transferred into a sterile microtube containing a few crystals of phenylthiourea to prevent melanization. Bacterial quantification was done by preparing 10-fold serial dilutions of hemolymph in PBS and inoculating the mixture onto THY agar plates supplemented with 5% sheep blood. Bacteria were enumerated by CFU counting after incubation at 37°C for 24 h, and results are presented as the number of CFU per milliliter of hemolymph.

## RESULTS

### Effect of pneumococcal RNase R on *Galleria mellonella* Infection

It was our main goal to evaluate the virulence potential of the *S. pneumoniae* RNase R mutant strain. In virulence studies, inoculation of the host with the same number of viable cells is essential. In order to evaluate its planktonic growth, we have used flow cytometry, allowing detection of live or death cells in the bacterial culture through the use of propidium iodide (PI) dye. The wild type and *rnr* mutant strain were grown in identical conditions, and the profile and number of live cells within the cultures were compared. The number of live cells at the same OD_600_ was identical for both strains (Figure S1A – Supplementary Material). The proportion of live and death cells were identical in the wild type and the Δ*rnr* strain, as judged by the PI ability to penetrate only the damaged, permeable membranes of non-viable cells. Indeed, the PI signal is detected in a low and identical number of cells for both wild type and RNase R mutant strain (Figure S1B – Supplementary Material). Our results indicate that deletion of RNase R has no impact in the number of cells during exponential growth.

To evaluate the virulence potential of the strain deficient in RNase R, we have compared the survival rates of *G. mellonella* infected with the wild type strain, the Δ*rnr* strain infection and the Δ*rnr* strain complemented with RNase R (Δ*rnr*+R). We have started by evaluating the pathogenicity level of the wild type strain by determining the dose of bacteria lethal for 50% of *G. mellonella*. For this, serial dilutions of the wild type were used to inoculate *G. mellonella*. Larvae survival and melanization was observed every 24 h until 72 h after inoculation. After 24 h of incubation, all the larvae infected with the wild type at 10^7^ bacteria/larva were death and presented strong melanization (Figure S2 – Supplementary Material). When 10^6^ bacteria/larva were used, 50% of larvae died and presented less melanization, while for doses below 10^6^ bacteria/larva all larvae survived even after 72 h of incubation. Thus, the lethal dose of the wild type bacteria necessary to kill 50% of larvae (LD_50_) is 5×10^6^ bacterial cells/larva, which is in agreement with the results previously obtained by [14].

To compare the survival rate of the larvae infected with the wild type and mutant strains, 5×10^6^ bacteria/larva (LD_50_) were thus used to inoculate *G. mellonella*. Survival of the larvae was evaluated as described above. As expected, the effects of infection by the wild-type strain were rapidly seen after the first 24 h of inoculation, with a reduction of about 50% of the initial larval population (Figure 1). After 72 h of incubation only 20% of the larvae survived. In contrast, deletion of RNase R increased by 30% the larval survival rate in the course of infection (Figure 1). Complementation of the mutant strain with RNase R expression *in trans* seemed even more lethal than the wild type after 24 h, but the wild-type survival rates were restored 72 h after infection (Figure 1).

These results demonstrate that the virulence potential of *S. pneumoniae* is attenuated in the absence of RNase R, establishing the importance of this ribonuclease in the capacity of this bacterium to cause infection.

**Figure 1.**
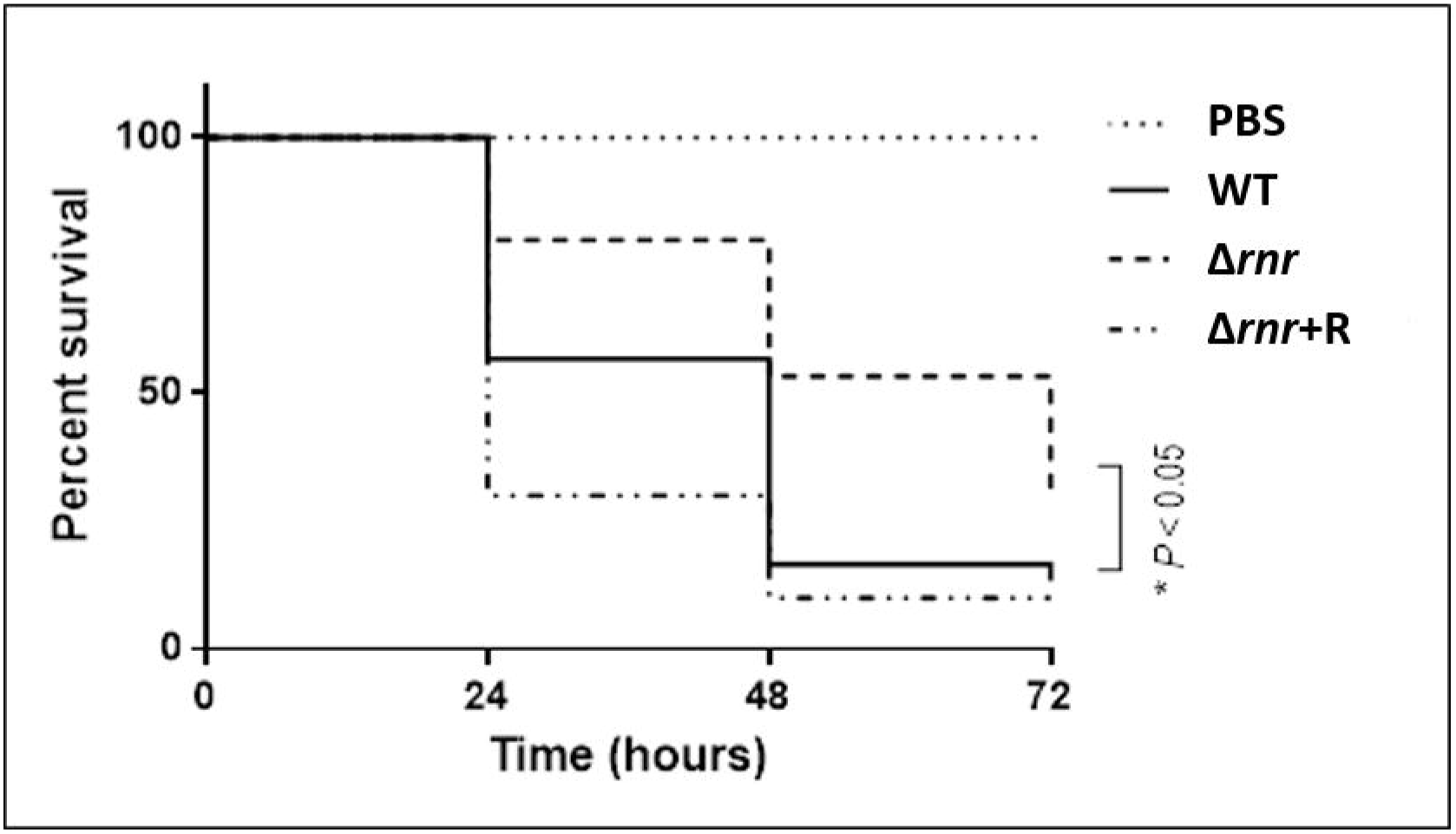
Survival of *G. mellonella* larvae after inoculation with *S. pneumoniae* wild type and RNase R mutant derivatives. Kaplan-Meier survival curves representing larvae infected with 5×10^6^ bacteria/larva. Larvae were infected with *S. pneumoniae* wild type (wt), *rnr* mutant (Δ*rnr*), Δ*rnr* expressing RNase R in trans (Δ*rnr*+R) and PBS (as a control) for 72 h. The results represent three independent experiments.

### RNase R Interference with *G. mellonella* Immune Response

Host defence peptides are a crucial part of insect innate immunity against invading pathogens, showing broad-spectrum microbicidal activity [25]. Therefore, it is expected that antimicrobial peptides are induced in *G. mellonella* upon *S. pneumoniae* infection as previously suggested by [14]. To test this, larvae were infected with this bacterium and the gene expression of four selected antimicrobial peptides was monitored at 1, 6, 12 and 18 h postinfection. The selected genes coded for the cysteine-rich antifungal peptide gallerimycin [26], a defensin called galliomycin [13], lysozyme [26], and an inducible metalloproteinase inhibitor (IMPI) [26]. Gene expression was determined by quantitative RT-PCR analysis of the total RNA extracted for each postinfection time point. Our results showed that infection with the wild-type TIGR4 strain led to upregulation of the immune-related peptides gallerimycin and galliomycin compared to the control larvae injected with PBS (Figure 2A), while no significant differences could be detected in the expression levels of IMP1 or lysozyme. We have then examined whether the absence of RNase R could be interfering with the activation of the immune system of the insect host, leading to the reduced virulence potential of the Δ*rnr* strain. To evaluate this, the same experiment was carried out using the wild type, the Δ*rnr*, and complemented (Δ*rnr*+R) strains. As shown in Figure 2B, the expression levels of all the tested immune-related peptides increased several fold within the larvae injected with the RNase R mutant strain, and the induction was generally more pronounced for the later times after injection. Surprisingly, even IMP1 and Lysozyme, which does not seem to be responsive to the wild type, were induced after infection with the Δ*rnr* strain. In all the cases, injection with the complemented strain partially reverted to the wild-type values, clearly indicating a role for RNase R in the activation of the insect immune response. This fact might be related with the lower virulence potential of the strain lacking RNase R.

**Figure 2.**
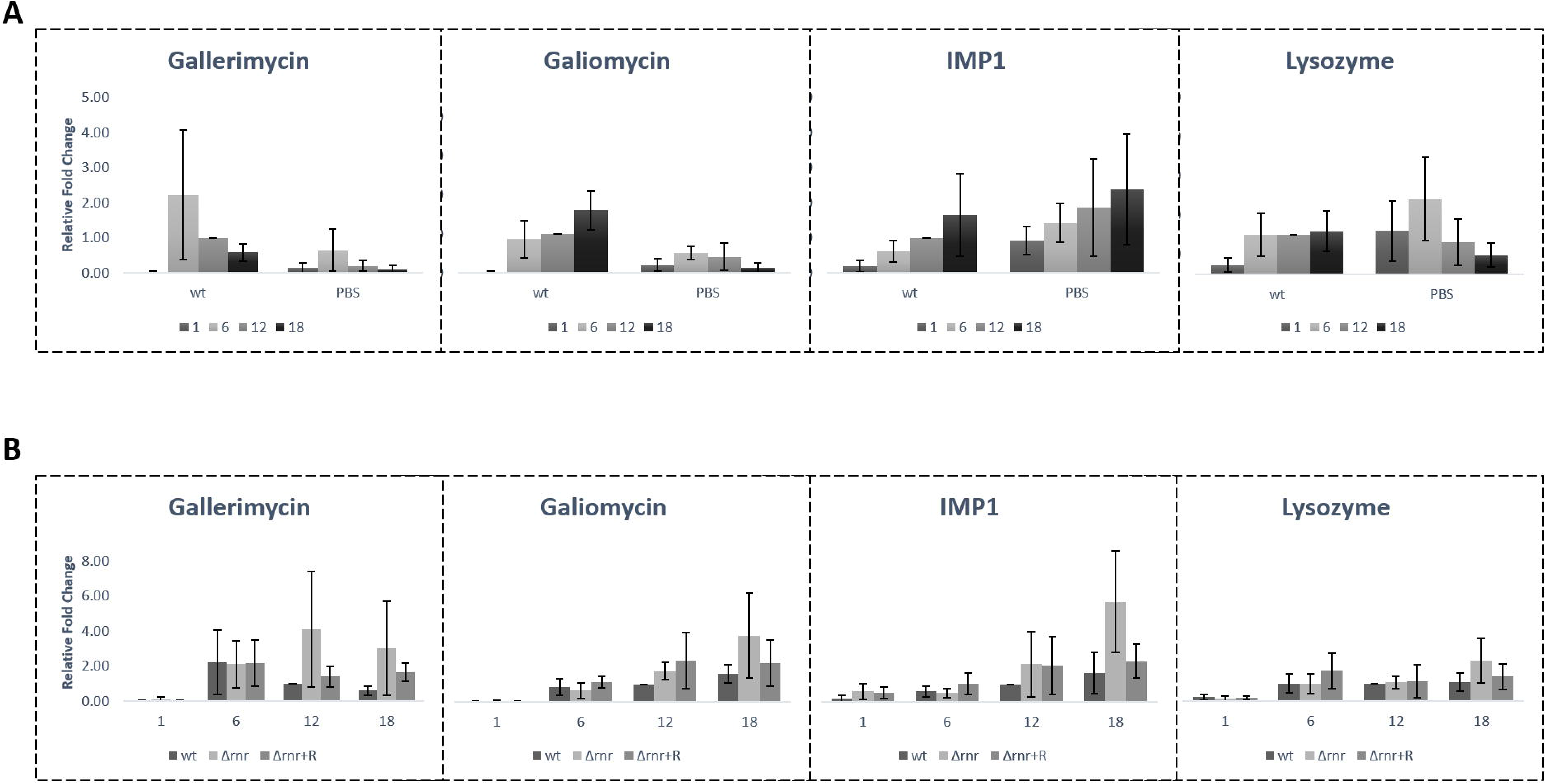
Immunogenic response of *G. mellonella* during infection with *S. pneumoniae* TIGR4 and derivatives. Expression levels of the immune-responsive genes of *G. mellonella* was measured at 1, 6, 12 and 18 h postinfection with bacteria/larva of *S. pneumoniae* TIGR4 wild type (wt), RNase R mutant (Δ*rnr*), Δ*rnr* expressing RNase R in trans (Δ*rnr*+R) and PBS (as a control). The transcriptional levels of gallerimycin, galliomycin, IMPI, and lysozyme were determined by quantitative RT-PCR analysis. **A)** Expression levels after infection with the wt strain compared to those in noninfected larvae injected with PBS. **B)** Comparison of the expression levels of the immune-responsive genes between the larvae infected with the wt and mutant strains. Results were normalized to the expression of the housekeeping gene actin and represent at least three independent experiments.

### RNase R Impact on the Bacterial Load in Hemolymph

High mortality is associated with bacterial proliferation within the host, while low mortality is associated with a bacterial clearance [14]. In order to evaluate the proliferation of *S. pneumoniae* within the insect hemocoel, we have determined the viable bacterial load within the hemolymph of larvae at three distinct time points in the course of infection. At 1, 6 and 18 h after injection of the larvae with *S. pneumoniae* wild-type, or RNase R mutant derivatives, hemolymph from three living larvae was collected and pooled, and the number of CFU was determined (Figure 3). Surprisingly, after 1 h the counts of the wild type have dropped down to near zero, and a sharp decrease was also observed for the strain complemented with RNase R, whereas the values of the RNase R mutant remained constant. Nonetheless at 6 h postinfection, both the wild type and the complemented strain start to recover and increasing numbers of bacteria could be detected in larval hemolymph to the point of 18 h postinfection. On the contrary the RNase R mutant suffered a severe decrease 6 h after infection and could not recover. At 18 h postinfection the bacterial load in hemolymph was significantly lower for the RNase R mutant than for the wild-type strain, which is indicative of a role for RNase R in the ability of *S. pneumoniae* to proliferate inside the host. The lower proliferation of the Δ*rnr* inside the larvae is in agreement with the higher survival of the larvae infected with this strain.

**Figure 3.**
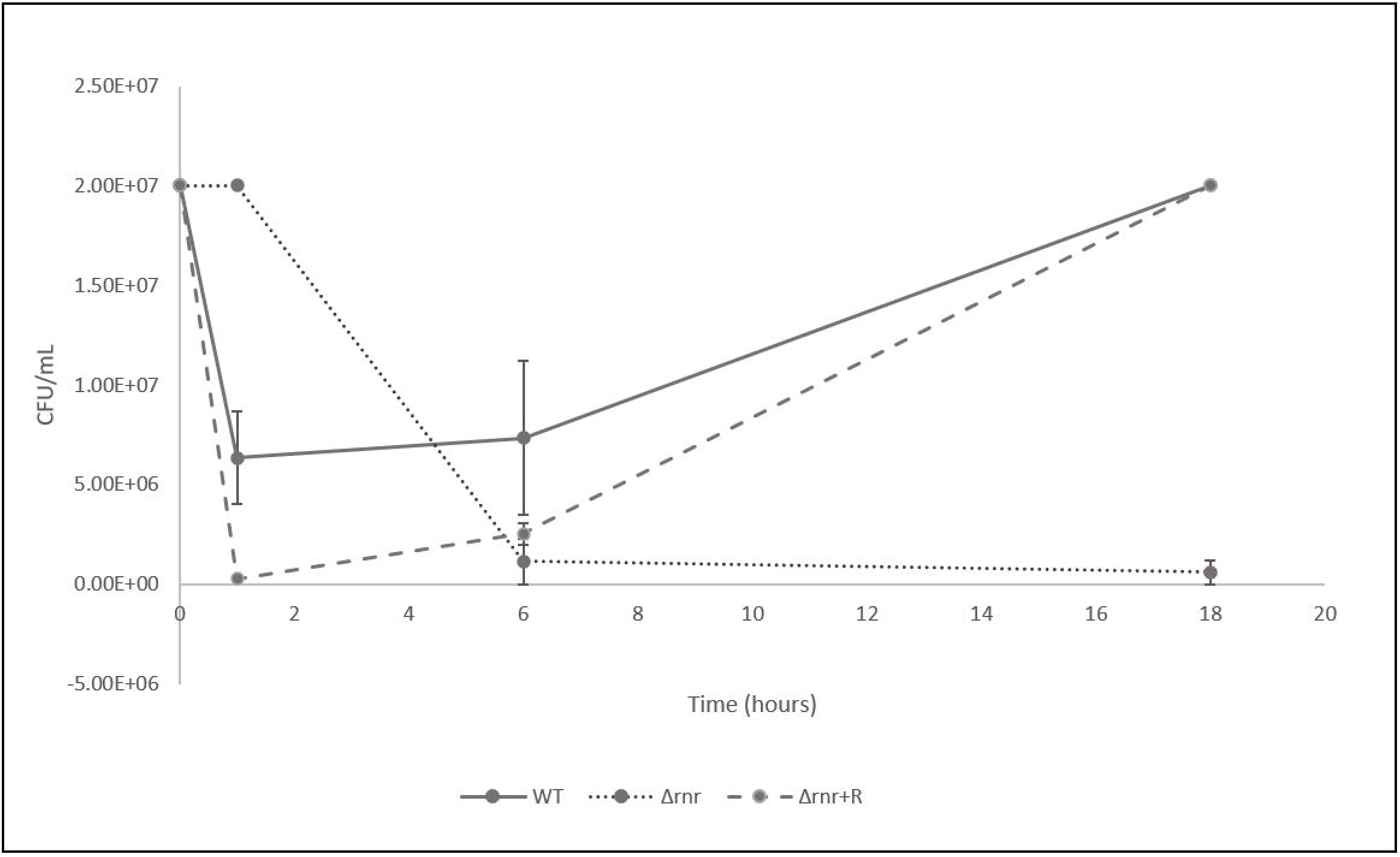
Hemolymph bacterial load in the course of *S. pneumoniae* infection. The viable bacterial load was determined in the hemolymph extracted from larvae infected with 5×10^6^ bacteria/larvae of the *S. pneumoniae* wild type (WT), RNase R mutant (Δ*rnr*), and complemented (Δ*rnr*+R) strains during a period within the time course of infection.

## DISCUSSION

Bacteria have developed a wide range of virulence mechanisms that are critical for successfully infect the host. A proper coordination of the bacterial cellular processes is essential for the progression of infection. RNase R is a stress response exoribonuclease, whose expression increases under several growth conditions [4, 27–30]. Involvement of this enzyme in virulence has been described in several pathogens [3] and its impact in protein synthesis and translation was recently shown in *S. pneumoniae* [2]. Therefore, in this work we have investigated whether RNase R would have a role in the pathogenicity of the pneumococcus. We show that the absence of RNase R leads to a decrease in *S. pneumoniae* virulence. Using *G. mellonella* as an infection model, our results demonstrate that 24 h after inoculating *G. mellonella* larvae with the wild type, only half of the larvae survived, while after infection with the Δ*rnr* strain, around 75% of the larvae survived during the same period. These differences were accentuated after 3 days, with a survival rate of only 10% after infection with the wild type, compared to 30% after inoculation with Δ*rnr*. Interestingly infection with the strain expression RNase R in trans even diminishes the survival rate. This is indicative of a direct role of RNase R in the pneumococcus pathogenicity, since in this strain RNase R levels are higher than in the wild type [31].

The Toll pathway, a crucial signalling cascade of the insect innate immune system, is activated by fungi and Gram positive bacteria leading to the synthesis of antimicrobial peptides [13, 32]. *G. mellonella* contains a notable antimicrobial peptide arsenal that is released into the hemolymph, where it attacks elements of the bacterial or fungal cell wall [21]. Among the antimicrobial peptides tested here, we found that infection with the wild type strain triggered the expression of gallerimycin and galliomycin. Infection with the deleted RNase R strain leaded to higher levels of these two immune-peptides, as compared to the wild type, and also induction of IMP1 and Lyzosyme expression. Hence, the absence of RNase R seems to induce a stronger humoral response by the larvae. This could be due to a higher immunogenicity of this strain but may also result from evasion of the immune response by the wild type, whereas the mutant would have lost this skill. Although the pneumococcus is considered a typical extracellular pathogen, its ability to proliferate within the host cells has recently been recognized [33–35]. Efficient clearance of pneumococci by macrophages and neutrophils was observed following intravenous inoculation of mice. However occasional intracellular proliferation of these sequestered bacteria provided a reservoir for subsequent dissemination of pneumococci into the bloodstream. The internalised bacteria evade clearance, undergo replication, then cause macrophage lysis and blood dissemination [33]. Endocytosis mediated Internalisation into cardiomyocytes followed by pneumococci replication within intracellular vacuoles was also reported [34]. In the lungs, uptaken pneumococci were proposed to persist intracellularly within dendritic cells and macrophages [36]. The accentuated decrease of the wild type and complemented mutant in the larvae hemolymph 1h after infection strongly resemblances the eclipse phase whereby pneumococci numbers rapidly decrease in the bloodstream after intravenous inoculation. Indeed, after the initial drop in bacterial numbers, a recover of both strains was observed with the respective load increasing until at least 18 h postinfection, resembling pneumococci blood dissemination after the eclipse. By contrast the bacterial load of the strain lacking RNase R, although higher 1 h after infection, constantly decreased and the strain could not recover. The other branch of innate immunity is cellular defence, mediated by *G. mellonella* hemocytes. They have some similarities with human macrophages and neutrophils, being able to phagocytise pathogens [13, 37]. Together our results are consistent with engulfment of pneumococci by the larval hemocytes, and are indicative of a role for RNase R in facilitating internalisation and/or replication inside the hemocytes. This would also explain the higher expression of the antimicrobial peptides in the larvae infected with the Δ*rnr*, which would thereby fail to be “hidden” inside the hemocytes. Our results indicate that the strains containing RNase R are able to evade the insect immune response, while in the absence of this enzyme the innate immune system is able to cope with the infection. RNase R might thus have an important role in the establishment of invasive disease, probably by affecting pneumococci phagocytosis. To invade and survive intracellularly, the pneumococcus utilizes a combination of virulence factors such as pneumolysin (PLY), pneumococcal surface protein A (PspA), pneumococcal adhesion and virulence protein B (PavB), exoglycosidases such as neuraminidase (NanA), the pilus-1 adhesin RrgA, pyruvate oxidase (SpxB), and metalloprotease (ZmpB) (reviewed by [35]). We are currently investigating the ability of the RNase R mutant to internalize and replicate inside *G. mellonella* larvae, as well as the influence of this ribonuclease in the expression of the above-mentioned pneumococcal virulence factors.

Overall, this work highlights the importance of RNase R in the ability of *S. pneumoniae* to establish infection, most probably by interfering with the pneumococcus interaction with the innate immune system of the host. The fact that the insect and human innate immune responses are very similar further raises the importance of this ribonuclease.

## Supporting information

Barria et al 2021 Supplementary Material

