## Supplementary material for "RNase R, a new virulence determinant of *Streptococcus pneumoniae*": Barria et al 2021 Supplementary Material

#### Figure S1

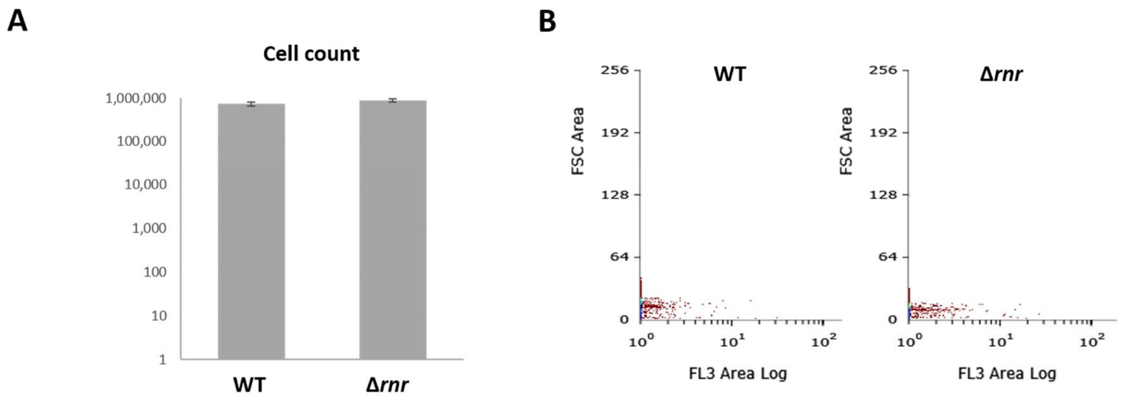


**Figure S1 – Flow cytometry analysis of *S. pneumoniae* wild type and Δ*rnr* cultures at exponential growth.** Analysis of *S. pneumoniae* wild type (WT) and *rnr* mutant (Δ*rnr*) cells growing in liquid media. **A)** Graphical representation of the number of live cells in each sample. **B)** Detection of propidium iodide signal. For all the experiments a low flow was used, and the percentage of live/dead cells was determined in 10 sec gated events. Unstained cells and dead cells from old cultures were used as negative and positive control, respectively.

### Figure S2

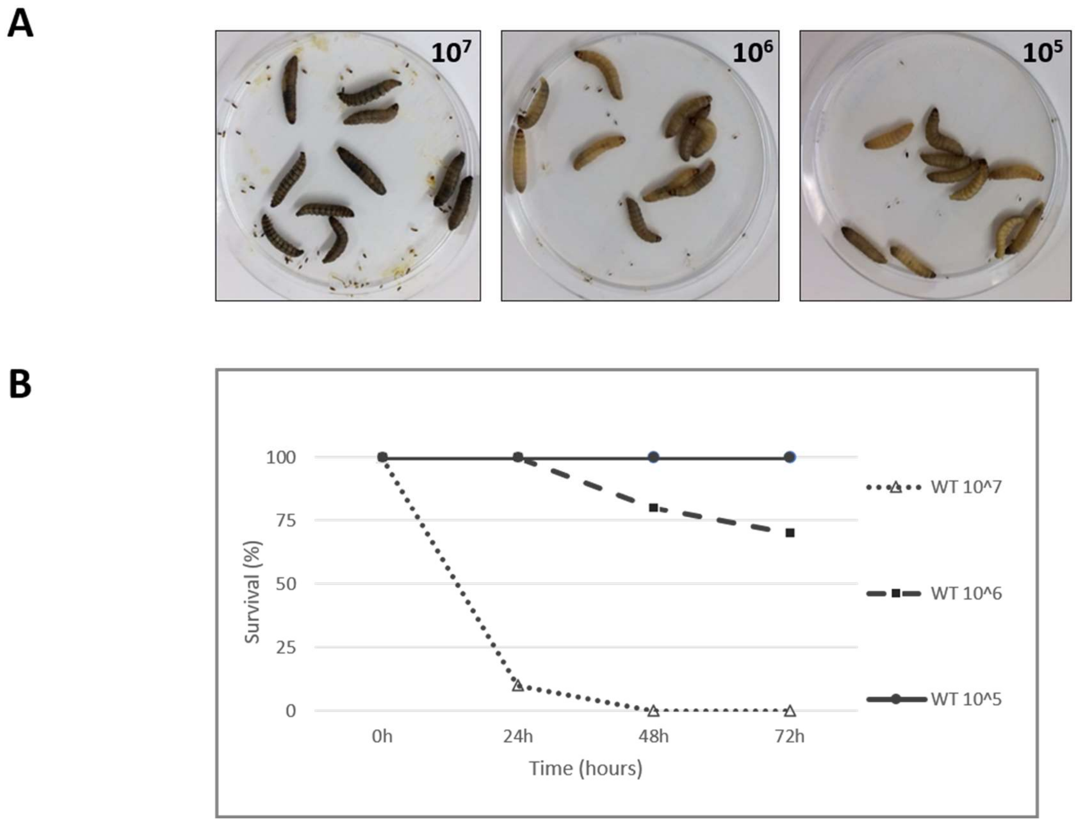


**Figure S2 – *G. mellonella* survival following infection with pneumococcal cells.** Larvae were inoculated with serial dilutions of *S. pneumoniae* wild type culture (number of bacterial cells is indicated on the top right corner of the corresponding photo, and on the right of the graph). Survival rate (**B**) and melanization (**A**) of the larvae after 24 h of incubation at 37°C are shown.

**Table S1** –Strains and plasmids used in this work.

| Strain/Plasmid | Relevant characteristics | Reference |
|---|---|---|
| **Bacteria** | | |
| ***S. pneumoniae*** | | |
| JNR7/87 (TIGR4) | | [1] |
| CMA607 | TIGR4 carrying pIL253 ($Ery^R$) | [2] |
| CMA611 | TIGR4 *rnr*⁻ (Δ*rnr*) ($Cm^R$) | [2, 3] |
| CMA604 | CMA611 carrying pIL253 ($Ery^R$) expressing RNase R (Δ*rnr*+R) ($Cm^R$) | [3] |
| CMA612 | CMA611 carrying pIL253 ($Ery^R$) | [4] |
| **Plasmids** | | |
| pIL253 | pAMβ1 derivative ($Ery^R$) | [2, 5] |
| pIL253-RNaseR | pIL253 carrying pneumococcal RNase R ($Ery^R$) | [3] |

$Ery^R$: Erytromycin resistant; $Cm^R$: Cloramphenicol resistant

**Table S2 –** Oligonucleotides used as primers in this work.

| Oligo name | Sequence 5' to 3' | Reference |
|---|---|---|
| P1RT (gallerimycin) | CGCAATATCATTGGCCTTCT | [6] |
| P2RT (gallerimycin) | CCTGCAGTTAGCAATGCAC | [6] |
| P1RT (IMPI) | AGATGGCTATGCAAGGGATG | [6] |
| P2RT (IMPI) | AGGACCTGTGCAGCATTTCT | [6] |
| P1RT (lysozyme) | TCCCAACTCTTGACCGACGA | [6] |
| P2RT (lysozyme) | AGTGGTTGCGCCATCCATAC | [6] |
| P1RT (actin) | ATCCTCACCCTGAAGTACCC | [6] |
| P2RT (actin) | CCACACGCAGCTCATTGTA | [6] |
| P1RT (galliomycin) | TCGTATCGTCACCGCAAAATG | [7] |
| P2RT (galliomycin) | GCCGCAATGACCACCTTTATA | [7] |
